# Global Transcriptome Characterization and Assembly of Thermophilic Ascomycete *Chaetomium thermophilum*

**DOI:** 10.1101/826354

**Authors:** Amit Singh, Géza Schermann, Sven Reislöhner, Nikola Kellner, Ed Hurt, Michael Brunner

## Abstract

A correct genome annotation is fundamental for research in the field of molecular and structural biology. The annotation of the reference genome *Chaetomium thermophilum* has been reported previously, but it is limited to open reading frames (ORFs) of genes and contains only a few noncoding transcripts. In this study, we identified and annotated by deep RNA sequencing full-length transcripts of *C.thermophilum.* We annotated 7044 coding genes and a large number of noncoding genes (n=4567). Astonishingly, 23% of the coding genes are alternatively spliced. We identified 679 novel coding genes and corrected the structural organization of more than 50% of the previously annotated genes. Furthermore, we substantially extended the Gene Ontology (GO) and Enzyme Commission (EC) lists, which provide comprehensive search tools for potential industrial applications and basic research. The identified novel transcripts and improved annotation will help understanding the gene regulatory landscape in *C.thermophilum*. The analysis pipeline developed here can be used to build transcriptome assemblies and identify coding and noncoding RNAs of other species. The R packages for gene and GO annotation database can be found under https://www.bzh.uni-heidelberg.de/brunner/Chaetomium_thermophilum.

## Introduction

The *Chaetomium thermophilum* is a thermophilic filamentous ascomycete, with the ability to grow at 50-52°C from degrading plant material (1). *C.thermophilum* produces, owned by it’s lingocellulolytic life style, different thermostable enzymes such as cellulase, xylanase, laccase, chitinases and proteases (2–6). The thermostability of these enzymes makes *C.thermophilum* a model organism of choice in various pharmaceutical and food processing industries. In the past decade *C.thermophilum* has attracted the attention for different applications such as starch degradation, hydrolysis of cellulose for bioethanol production as well as other applications requiring enzymatic activities at higher temperatures (6–14). Additionally, many *C.thermophilum* proteins and complex protein assemblies have been reported in highest resolutions in various crystallisation and cryo-electron microscopy studies, which improved our understanding of the structural organization and function of protein complexes. These include the Crm1 export factor, the splicing factor Cwc27, mRNA export factor Mex67-Mtr2, the FACT complex, the eukaryotic RAC chaperone, the nuclear pore Nsp1-channel complex, and the 90S pre-ribosomal complex (15–22). The genome of *C.thermophilum* was first reported by (23) and afterwards substantially improved in its annotation by (24). It has a size of 28.3 Mb and assembled into 20 scaffolds containing 7165 protein coding and 387 noncoding transcripts. Despite of recent advances in sequencing technologies (25–28) the *C.thermophilum* genome has not been substantially upgraded. The reported *C.thermophilum* genome lacks proper annotation of untranslated regions (UTRs), and the majority of intron-exon structures are computationally predicted rather than experimentally determined. Given the substantial increase in the number of genomic studies on *C.thermophilum* a comprehensive genome annotation will be helpful for further functional, structural, proteomic, genomic and transcriptomic analyses. In this study, we present an improved annotation of the *C.thermophilum* genome based on deep RNA sequencing and establish pipeline tools for the analysis of sequencing data. Our annotation identified 7044 expressed protein-coding genes and 4567 long noncoding RNAs (lncRNAs). Moreover, our pipeline detected UTRs and intron-exon boundaries as well as transcript isoforms. Sequence homology studies revealed that *C.thermophilum* and *Thermothelomyces thermophila* share close sequence similarity of coding transcripts. The downstream analysis of genomic and transcriptomic sequence data is often used to predict the function of genes, identification of biomarkers, grouping and classifying gene expression patterns. Therefore, we present an extended Gene Ontology (GO) and Enzyme Commission (EC) numbers that are associated with protein coding genes of *C.thermophilum*.

## MATERIALS AND METHODS

### RNA Isolation and sequencing

The *C.thermophilum* strain was received from DSMZ, Braunschweig, Germany (No. 1495). The mycelium scraped off a freshly grown agar plate were grown in 250 ml baffled Erlenmeyer flasks at 52°C for 6 h. Mycelium was grown in modified CCM medium as described in (29): 0.5 g NaCl, 0.65 g K_2_HPO_4_·3H_2_O, 0.5 g MgSO_4_·7H_2_O, 0.01 g Fe(III)-sulfatehydrate, 8 g D-glucose and 1 g each of peptone and yeast extract per liter H_2_O, pH 7.0. *C.thermophilum* mycelia were harvested through a sieve and grounded to a fine powder in liquid nitrogen. About 100 mg mycelia from three independent biological replicates were used directly for total RNA-extraction using the SV total RNA isolation system (Promega). The libraries were prepared with the NEBNext Ultra II Directional RNA Preparation Kit for Illumina in combination with NEBNext Poly A selection Module, plus the NEBNext Multiplex Oligos for Illumina, and sequencing was performed by CellNetworks Deep Sequencing Core Facility (Heidelberg, Germany) on Illumina NextSeq 500 platform (single-end).

### RNA sequencing data analysis

The data quality assessment of raw sequence data was performed by FastQC (Version: FastQC 0.11.5, (http://www.bioinformatics. babraham.ac.uk/projects/fastqc/). No samples were discarded from the analysis. *C.thermophilum* reference genome and gene annotation files were downloaded from ENSEMBLE (https://fungi.ensembl.org) and a pipeline was developed to annotate UTR region and to identify the putative novel transcripts (Figure −1A). The raw reads were mapped to *C.thermophilum* genome using HISAT2 with the following parameters (Version: 2.1.0; [hisat2 -p 8 -x -max-intronlen 2000 -dta-U])(30). The mapped reads from HISAT2 for each sample were assembled separately using StringTie with parameter settings (Version: 1.3.3b; [stringtie -o -m 50 -p 8 -j 3 -c 5 -g 15])(31). The multiple transcript assembly files from the different samples were used together to produce a distinctive transcriptome set using gffcompare with parameter settings (Version: v0.10.1; [gffcompare-merge -K -o gffcomp -i]) (25). Based on the previous assembly results, transcripts shorter than 200 nt were excluded to identify transcripts from the merged transcript assembly. According to the gffcompare class codes “i”, “u”,”y” and “x” are considered as novel transcriptional loci. The coding potential calculator (CPC2) was used to evaluate the coding potential of all transcripts (32).

**Figure-1:**
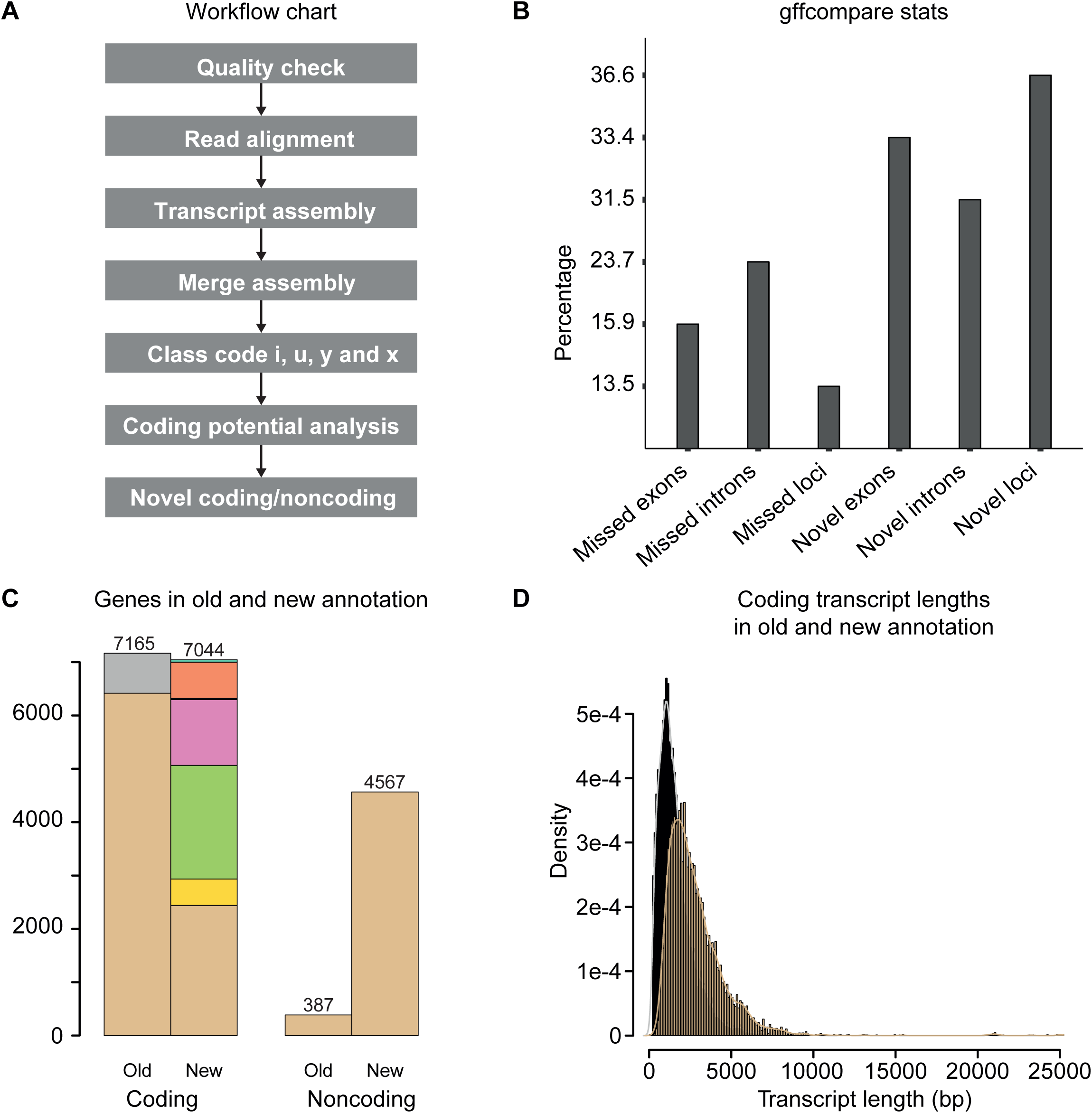
Transcriptome-based annotation of the genome of *C.thermophilum*. A) Systematic overview of the analysis pipeline. B) Identification of the novel intron and exon loci in percentage from newly annotated genome. C) Stack bar plot of genes of the old and new annotation. The stacked grey bar for the coding genes in the old annotation represents genes that were not expressed in our conditions. In the stacked bar of the new annotation, the colors represent differences compared to the previous annotation. The individual colors indicate for the following features: Beige; genes without change in the intron structure (n=2440). Yellow: genes where at least one isoform has a novel splicing variant (n=493). Green: genes where all transcripts have at least one intron-exon junction different than in the previous annotation (n=2132). Pink; genes where all junctions are different (n=1234). Blue; genes that are flipped (opposite strand with same or similar splice junctions) (n=19). Orange; completely novel genes (n=679). On the top the aquamarine represents the uncategorized genes (n=47). D) Histograms representing the coding transcript lengths from the new *C.thermophilum* genome annotation (beige) compared to the length of the previous annotation of ORFs (black). A detailed description of the methods is given in the respective section of the main text.

### Sequence conservation and GO annotation

The sequence conservation analysis was performed using dc-mega BLAST (Version: 2.7.1+) (33). All coding and noncoding transcripts including the identified novel transcripts sequences were used for this analysis. *Sordaria macrospora* (NCBI taxid-5147), *Neurospora crassa* (NCBI taxid-5141), *Aspergillus niger* (NCBI taxid-5061), *Saccharomyces cerevisiae* (NCBI taxid-4932), *Takifugu rubripes* (NCBI taxid-31033) and *Thermothelomyces thermophila* (NCBI taxid-78579) were chosen to study the sequence similarity analysis using BLAST (E value,1e^-3^). phyloT (https://phylot.biobyte.de) was used for the construction and visualization of phylogenetic tree of the above mentioned species. Additionally, functional annotation of *C.thermophilum* transcripts were analyzed using Blast2GO [Version 5.1.1] (34) as described in the manual. The annotated GO terms from *Thermothelomyces thermophila, Neurospora crassa* and *Sordaria macrospora* were used as an input for the Blast2GO analysis, based on local blastx. Enzyme Commission numbers were obtained using the same method. The data visualization was carried out using R (Version 3.3.3; http://www.R-project.org/) (35).

### Isoform annotation

To create the isoform annotation, the 9755 coding transcripts were analyzed. The longest non-reverse ORF, in search order of [blastx-hit -frame1 -frame2 -frame3], using ATG only as start codon and obligatory stop codon was obtained from Blast2GO. Sequences were grouped by the transcription loci tags pyfaidx python package (36). All groups were aligned [-output=aln] and similarities were calculated [-other_pg seq_reformat -output sim] using T-Coffee software (37). Pairwise similarity scores formed two distinct groups. Based on this, the score cutoff level was set at 49 for designating isoforms. All low scoring accepted hits (score 49-60) were manually checked and visualized for correctness. The accepted protein sequence pairs were merged into isoform groups by connectivity calculation in R. Noncoding transcripts were grouped by overlapping features.

## RESULTS

### Transcripts reassembly and Identification of novel transcripts

The single end RNA seq data of *C.thermophilum* was obtained in triplicates with a length of 85bp. We performed short read gapped alignment using HISAT2 (30) and recovered more than 95% mapped reads as shown in Table-1. Further, we used StringTie (31) to de novo assemble all three samples separately. The assembled transcript files from these three samples were merged into a combined set of transcripts using the gffcompare utility provided by Cufflinks. After manual curation (17 transcripts) a total of 15363 reliable transcripts were obtained after filtering the transcript length (>200nt) (Figure-1C). In total, the transcripts correspond to 7044 coding genes, represented by 9772 transcripts and transcript isoforms, and 4567 noncoding genes, represented by 5591 transcripts and transcript isoforms. Gffcompare statistics on the comparison of the found transcripts compared to the previous annotation are shown in Figure-1B, and the transcripts annotated class codes are listed in Table-2, Figure-1D represents the increased lengths of the newly annotated transcripts compared to the earlier annotation, which did not include UTRs. Transcripts annotated to gffcompare classes u (no overlap, n = 2744) and x (opposite strand, n = 1754), as well as i (contained in reference intron, n = 28) and y (contains a reference gene within intron, 5 transcripts) are called as a novel transcript in our analysis. By these criteria, we identified 679 novel coding genes represented by 892 transcripts and isoforms as well as 2878 novel noncoding genes represented by 3639 transcripts and isoforms. 749 genes that were classified as a coding in the previous annotation were not detected/expressed under our experimental condition. These non-expressed putative genes are less conserved in related fungi (below 40%), suggesting that this group may contain falsely annotated genes. The total number of newly identified coding and noncoding transcripts is shown in Figure-1C. Finally, we created a gene annotation database containing all above information in TxDb framework in R package for *C.thermophilum* (38). The gene annotation file in gtf format can be found in supplementary file-1.

### Annotation of the novel noncoding RNAs in *C.thermophilum*

We identified 5591 noncoding RNA transcripts based on CPC2 analysis. These include highly expressed contaminant RNA species such as ribosomal RNAs, t-RNAs, snoRNAs, RNase RNAs and snRNAs. The remaining non-coding transcripts were classified as intronic, intregenic, sense overlap with coding gene and antisense based on association with annotated protein-coding genes as shown in Figure-2D. In total we identified 2188 lincRNA genes as intregenic, 1949 antisense and 530 genes overlapping in sense direction. 166 genes are in both sense and antisense, these could represent truncated mRNAs or functional RNAs involved in gene regulation as shown in Figure-2C.

**Figure-2:**
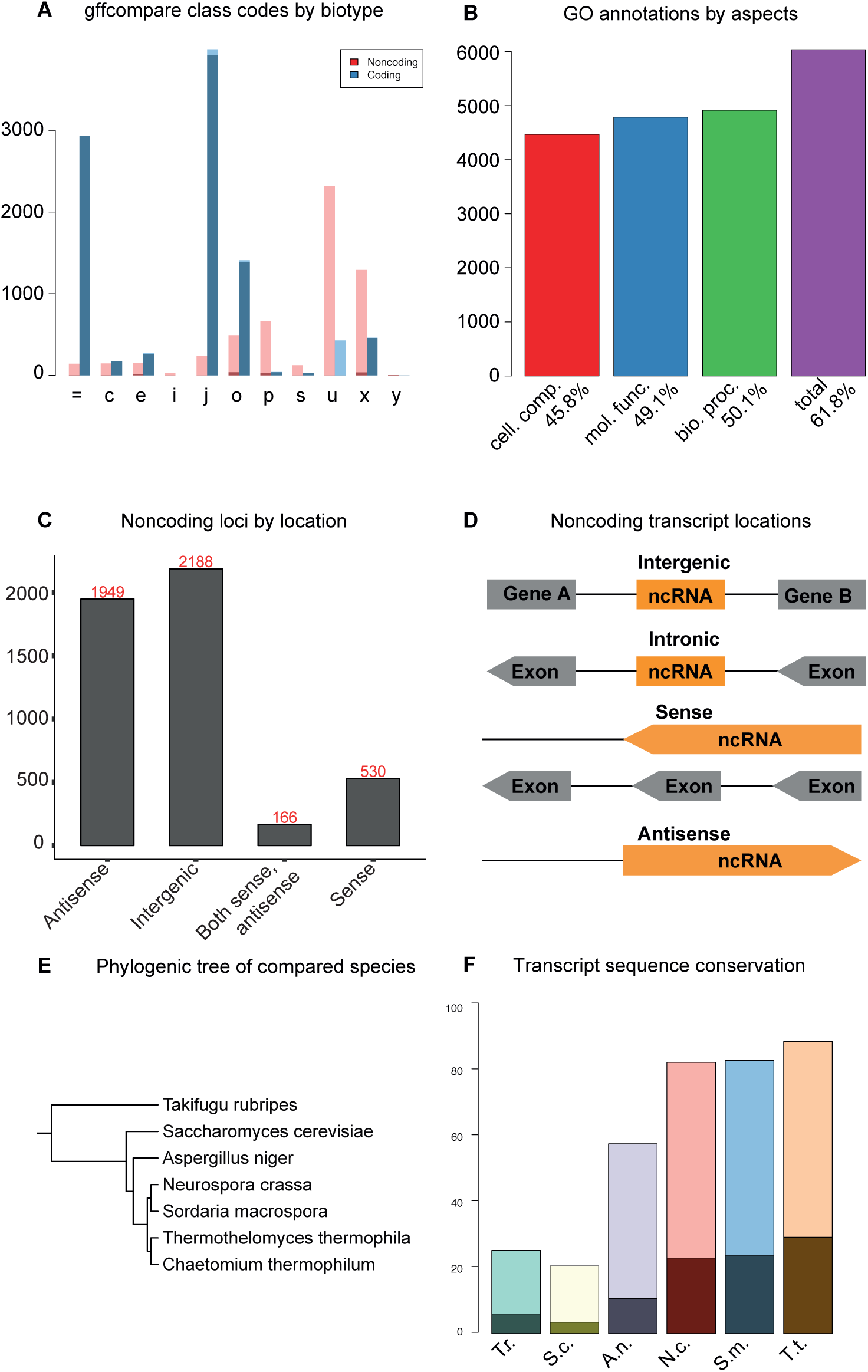
A) Transcript assemblies and gffcompare class codes. Stack bar plot represents the gffcompare class codes containing coding and noncoding transcripts. Darker shading represents the part where the corresponding transcript in the old annotation has the same biotype. B) GO annotations associated with genes, representing biological process (BP), molecular function (MF) and cellular compartment (CC). C) Bar plot representing different types of noncoding genes. D) Graphical notation of different types of noncoding transcripts by location. E) Phylogenic tree of 6 compared species. F) Stack bar plot representing the sequence similarity of transcripts of 6 compared species. Darker shade for the non-coding transcripts, light shade for the coding.

### Sequence conservation and functional annotation

A phylogenetic tree including 6 other species indicates that *C.thermophilum* is closely related to *Thermothelomyces thermophila* and more distantly to *Neurospora crassa* and *Sordaria macrospora* (Figure-2F). Local dc-megaBLAST similarity search was carried out for the newly annotated coding, anti-sense and lincRNAs of *C.thermophilum*. We found that 90% of the coding and 45% of the non-coding sequences resulted in a significant hit with *Thermothelomyces thermophila* (Figure-2E). We also observed that both *Sordaria macrospora* and *Neurospora crassa* appear to be sharing 80% coding and 25% noncoding sequence similarity (Figure-2E). The similarity search against the other three species revealed lower sequence similarities as shown in Figure-2A. The gene ontology (GO) facilitates the functional characterization of genes; therefore, Blast2GO was used to associate the transcripts with a functional annotation. We found altogether 4283 GO terms, 1428 belong to Molecular Function (MF), 2140 to Biological Process (BP), and 715 to Cellular Component (CC)(Figure-2B). With these, we could annotate 4336 coding genes. All GO terms are listed in supplementary file-2 as well as top 10 GO-slim terms for each category (MF, CC, BP) are shown in supplementary Figure-1. Further, we created a GO annotation R package for *C.thermophilum* using function makeOrgPackage (38). To facilitate functional gene finding for industrial applications, we also retrieved Enzyme Commission numbers (E.C) from the Blast2GO analysis. We could associate 1366 coding genes with 643 E.C. numbers. The main EC classes distribution is shown in Figure-3.

**Figure-3:**
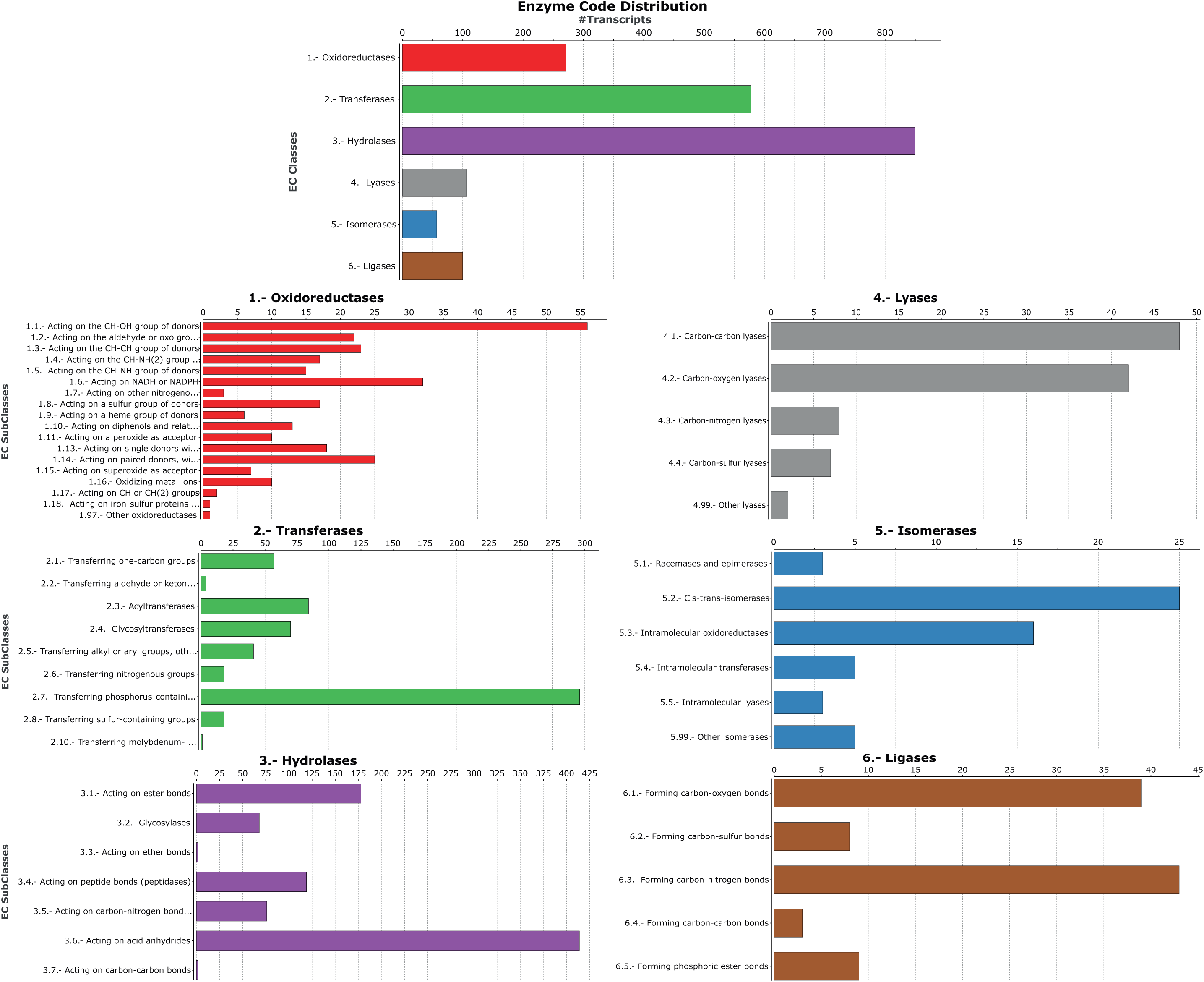
Bar plot enzyme class code distribution with transcripts. 6 different enzyme classes are distributed throughout the transcripts. Each bar plot (1-6) containing top 5 enzymes with specific enzyme class code.

## DISCUSSION

*C.thermophilum* is an economically important species in pharmaceutical and food processing industries, and a useful model organism in basic research, particularly in the field of ribosome biogenesis (12–14, 22, 39, 40). In this study an analysis pipeline was developed to characterize the transcriptome of *C.thermophilum*. The main aim of the present study was to provide a comprehensive gene annotation, containing high fidelity coding and noncoding transcripts and isoforms with both 3’ and 5’ UTRs. We evaluated all transcript class codes from our transcriptome assembly and compared it with the previous annotation. We observed substantial discrepancies with the previous annotation of *C.thermophilum*, as only 2935 coding and 143 noncoding transcripts displayed a complete intron match (class code “=“). The majority of coding transcripts showed at least one intron mismatch with the old annotation (class code “j”). We observed that 749 coding and 254 noncoding transcripts of the previous annotation were not expressed (no overlaps with any identified transcript) in our analysis. 104 of these genes were potentially expressed in other growth conditions (> 100 read counts, unpublished RNA sequencing data), while the remaining genes were not expressed. Furthermore, most of these genes had no homology in related species, suggesting that they might have been wrongly annotated. These unexpressed transcripts are listed in supplementary files-4.

Our transcript assembly revealed that 1640 genes express at least 2-3 transcript isoforms (4368 transcripts, 44.7%), indicating rather complex alternative splicing in *C.thermophilum*. A hand full of genes may express even higher numbers of isoforms. However, due to theoretical limitations in the analysis of the single end sequencing data such potentially complex isoforms cannot be reliably predicted. Here we present all potential transcript isoforms, and the corresponding predicted protein sequences are listed in supplementary file-3. Our analysis also revealed a surprisingly high number of noncoding transcripts, both lincRNA and antisense RNAs. Taken together with the alternative splicing of coding genes our data suggests an astonishing complexity of the genome architecture and transcriptional network of *C.thermophilum*. Such complexity is usually found in higher eukaryotes where it is supposed to permit complex life style. Comparative analysis uncovered that *Thermothelomyces thermophila* shows 90% genome sequence similarity with *C.thermophilum* as shown in Figure-2E). The high sequence similarity of *Thermothelomyces thermophila* and *C.thermophilum* is likely to reflect related functions, potentially associated with their thermophilic lifestyle. Functional annotation through Gene Ontology (GO) associations facilitates the interpretation of genomic and transcriptomic sequence data. Functional annotation achieved by the integration of several databases such as KEGG (43), UniProt (44), InterPro(45), Pfam(46), NCBI (47), SEED (48), ConsensusPathDB (49) and Reactome (50) may help addressing this question. Our Blast2GO analysis substantially expands the GO term annotations in *C.thermophilum*. Hence, our annotation and transcript assembly opens opportunities for the systematic analysis of *C.thermophilum*. An example would be a known protein fragment of a laccase enzyme from *C.thermophilum* (Uniprot ID: Q692I0) that was so far not annotated in the genome and the full protein sequence was unknown. We found the gene expressed in our conditions and have annotated the complete transcript and protein sequence (Gene001724). *C.thermophilum* genes responsible for the formation conidia are another example for an extended functional annotation. The reproduction cycle of *C.thermophilum* is not well understood, and previously only a single gene had been associated with the formation of asexual spores (conidia). Our GO term annotation revealed that several additional genes, such as Gene000283; Gene003743; Gene006754; Gene006902, are associated with the conidiation process (GO:0048315, GO:0030437, GO:0030435).

Our analysis is solely based on a single growth condition with single-end RNA seq data. Hence, different growth conditions, as well as paired-end sequencing data could further improve the gene annotation. RNA degradation seems to be a general problem in *C.thermophilum*. Therefore, improvement of the RNA isolation procedure could decrease the number of truncated transcripts. In summary, we detected a large number of novel coding and noncoding transcripts and discovered a high diversity of alternative splicing in *C.thermophilum*. Thus, our study provides useful resources for functional genomics and proteomics research on *C.thermophilum* and facilitates the analysis of biological and biochemical processes. Moreover, our analysis pipeline can be used for the genome annotation of other organisms.

## AVAILABILITY

The GO Term and gene annotation R packages of *C.thermophilum* are available in the following link. The gene annotation R package can be found in https://www.bzh.uni-heidelberg.de/brunner/Chaetomium_thermophilum/TxDb.Chaetomium.ct39.knownGene_1.0.0.tar.gz and GO annotation can be found in https://www.bzh.uniheidelberg.de/brunner/Chaetomium_thermophilum/org.Cthermophilum.eg.db_1.0.0.tar.gz

## ACCESSION NUMBERS

The RNA seq data are deposited in the Gene Expression Omnibus database under accession no GSE116834.

## SUPPLEMENTARY DATA

We provide the following files for both scientific and industrial applications.

1. The C.thermophilum_new.gtf file containing the essential attributes/annotation features that can be used further in genomics and transcriptomics data analysis of *C.thermophilum*
2. The C.thermophilum.xlsx file containing all genes associated with GO annotation and E.C. numbers.
3. The C.thermophilum protein.fasta file contains all the translated protein sequences, ORF with both start and stop codons in the forward strand, with at least 50 AA lengths.
4. Missing.genes.xlsx contains the 749 coding genes from the old annotations where we found no expression in excel format.

## ACKNOWLEDGEMENTS

We thank David Ibberson at the CellNetworks Deep Sequencing Core Facility (Heidelberg, Germany) for performing library preparation and HiSeq sequencing. AS thanks Prof. Dr. Thomas Höfer and Dr. Congxin Li (Bioqunat, Heidelberg, Germany) for allowing processing the sequencing data in their computer cluster. As thanks to PD Dr. Jochen Baßler for fruitful discussion.

## TABLE AND FIGURES LEGENDS

Table-1: RNA-seq data alignment results for reads of different samples. Table-2: Different classes of assembled transcripts

**Supplementary Figure-1:**
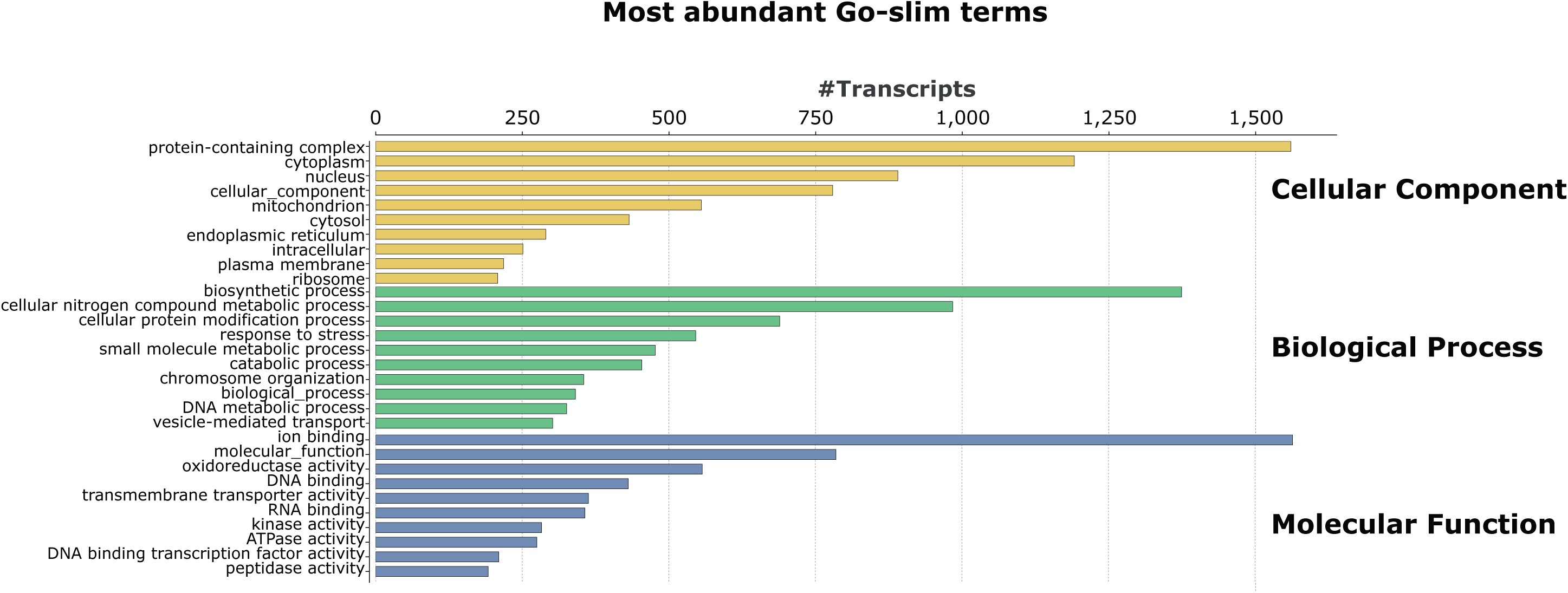
Bar plot of top abundant GO slim terms obtained from Blast2GO analysis. Y-axis represents the GO term and x-axis represents the total number of transcripts are involved in particular term. Different color code such as light-yellow corresponds to the cellular component and light green to biological process and light blue to molecular functions.

