## Supplementary material for "Global Transcriptome Characterization and Assembly of Thermophilic Ascomycete *Chaetomium thermophilum*": Table 1

Table- 1: RNA-seq data alignment results for reads of different samples.

| Sample name | Input read | overall alignment rate |
| --- | --- | --- |
| G1 | 69350606 | 95.94% |
| G2 | 73230226 | 95.87% |
| G3 | 59976502 | 95.83% |
