## Supplementary material for "Global Transcriptome Characterization and Assembly of Thermophilic Ascomycete *Chaetomium thermophilum*": Table 2

Table- 2: Different classes of assembled transcripts

| Class code | Description | Total annotation |
| --- | --- | --- |
| = | Complete match of intron chain | 3078 |
| c | Contained in reference (and intron isoform compatible) | 323 |
| k | containment of reference (reverse containment) | 0 |
| j | At least one splice junction match | 4234 |
| e | At Single exon, overlapping intron a possibly pre-mRNA fragment (un spliced intron) | 420 |
| o | Other same strand overlap with reference exons | 1896 |
| s | Intron match on the opposite strand (likely a mapping error) | 158 |
| x | Exonic overlap on the opposite strand (like ,’o’ or ’e’ but on the opposite strand) | 1754 |
| i | fully contained in a reference intron | 28 |
| y | Contains a reference within is intron(s) | 5 |
| p | Possible polymerase run-on (no actual overlap) | 706 |
| r | repeat (at least 50% bases soft masked) | 0 |
| u | none of the above (unknown,intergenic) | 2744 |
